# High-resolution MRI Guided Whole Mouse Brain Cell Type Atlas using Deep Learning

**DOI:** 10.1101/2025.11.25.690475

**Authors:** Xinyue Han, Rui Hu, Zhuoheng Liu, Jie Chen, Mubashir Jafry, Haoyu Song, Yi Zhao, Mingquan Lin, Leonard E. White, G. Allan Johnson, Nian Wang

## Abstract

Cell types represent groupings of cells defined by shared anatomical and functional properties. Traditional mouse brain cell atlases rely heavily on single-cell sequencing, which provides valuable molecular detail but lacks whole-brain, isotropic resolution. Diffusion MRI (dMRI) offers a complementary approach for probing cytoarchitecture and myeloarchitecture, with quantitative metrics increasingly used as biomarkers of brain development and neurodegenerative disorders. Although prior work has linked dMRI metrics with gene expression, the capacity of dMRI to directly predict cell types remains unclear. Here, we present a deep learning framework that integrates high-resolution dMRI with three-dimensional light-sheet microscopy (LSM) registered to the Allen Mouse Brain Common Coordinate Framework (CCFv3). We investigated correlations between dMRI and spatial transcriptomics-derived cell types and generated a whole-brain cell type atlas at 10 µm isotropic resolution. Together, these results establish an efficient, high-resolution strategy for brain cell atlas generation and underscore the potential of advanced imaging to illuminate cellular mechanisms of the brain.

## 1. Introduction

Cell types are the basic structural and functional units of organisms. In the mammalian brain, cell types typically have a high degree of diversity and specialization. Studies^1,2^ have shown that cell types are highly diverse in the midbrain, hindbrain, hypothalamus, and cortex. Yao et al.^3^ found that cell types have a high degree of transcriptomic identity and spatial specificity. More importantly, brain cell types also have a hierarchical feature in terms of taxonomies, including (from the highest to the lowest): (1) neuronal and non-neuronal (glial) classes, (2) cell classes by neurotransmitter type and major brain regions, and (3) cell subclasses within each cell class.^3,4^ Characterization of brain cell types is important, as it will facilitate our understanding of the development, evolution, and pathology of the brain^5^. One of the most widely used approaches is single-cell RNA-sequencing (scRNA-seq), which can profile the expression of thousands of genes per cell. By combining scRNA-seq and spatial transcriptomics, comprehensive cellular and molecular brain atlases can reveal cell types and their epigenomic information in their native tissue context. Brain Initiative Cell Census Network (BICCN) is developing several mouse brain atlases, including transcriptomic and spatial atlas,^1,3,6^ epigenetic landscape and gene regulatory elements.^7,8^ Although many of the aforementioned atlases have been integrated into the Allen Mouse Brain Common Coordinate Framework (CCFv3),^9^ limited efforts have been made to enable predictive interference of one modality from the other.^10^ As cell atlas research is fundamental for guiding future therapeutic innovations for brain disorders,^11^ cross-modal prediction becomes increasingly essential for validating the consistency and alignment of cell type identities across modalities.

Novel imaging techniques, such as magnetic resonance imaging (MRI), are one alternative to scRNA-seq for characterizing brain cell types. Among various MRI techniques, diffusion MRI (dMRI) provides a unique approach to measure the diffusion of water molecules in tissue and has been demonstrated to reveal tissue microstructure and cytoarchitecture.^12^ For example, the apparent diffusion coefficient (ADC) derived from diffusion-weighted imaging (DWI) is highly correlated with cellular size changes during acute brain ischemia.^13^ By using different statistical and biophysical models, dMRI can detect specific cellular changes. Diffusion tensor imaging (DTI) applies the diffusion tensor model to extract the anisotropy and diffusivity metrics. The degree of diffusion anisotropy during brain development reflects the migration and organization of glial cells and neurons^14^ and the myelination process^15^. Diffusion kurtosis imaging (DKI) quantifies the non-Gaussian distribution of water diffusion and has a high sensitivity for pathological changes in neuronal tissues.^16^ Neurite orientation dispersion and density imaging (NODDI) employs a biophysical model to extract the neurite density index (NDI) and orientation dispersion index (ODI) of tissue. NODDI has better specificity for certain tissue properties, such as its superior sensitivity to tissue-specific white-matter tract damage in Alzheimer’s Disease.^17^ Soma and Neurite Density Imaging (SANDI) is more specific to cellular and neurite densities, as the tissue component model explicitly includes the soma of any brain cell type, enabling the estimation of apparent soma density and size.^18^ Current studies have investigated the association of dMRI parameters with cell type-specific spatial transcriptomics. De Santis et al.^19^ found that mean diffusivity (MD) is sensitive to microglia density. Patel et al.^20^ found that fractional anisotropy (FA) is associated with CA1 pyramidal cells. Shen et al.^21^ found that FA is correlated with myelination and oligodendrocytes, and the diffusivity indices are correlated with neurons and their connecting compounds. However, whether dMRI can predict brain cell types is still unknown.^22,23^ Investigating the predictive performance of dMRI in brain cell types will enhance its applications in neuroscience and deepen our understanding of the brain across different scales.

In this study, we integrated high-resolution diffusion MRI and high-resolution 3D light-sheet microscopy (LSM) into CCFv3 space, and built a deep learning-based whole-brain cell type prediction model termed MFNet to predict mouse whole-brain cell atlas from Allen Brain Cell (ABC) Atlas (Fig. 1). Firstly, we segmented neuronal cells from 3D NeuN images (Figure 1A), acquired high-resolution (45 µm isotropic) *ex vivo* mouse brain diffusion MRI images, and fit statistical and biophysical diffusion models including DTI, DKI, NODDI, and SANDI (Figure 1B). Secondly, we registered the dMRI images and the 3D LSM to the CCFv3 space. Thirdly, according to the hierarchical and region-specific properties of the cell type dataset^3^ (Figure 1C), we trained a Multi-Diffusion-MRI Fusion Network prediction model (MFNet) on neuronal cell type with a three-level taxonomical classification feature and a co-localization-based brain division stratification strategy (Figure 1D). We validated the prediction accuracy by using leave-p-groups-out cross-validation. Furthermore, we examined the correlation between dMRI parameters and spatial transcriptomics of neuron-type marker genes (Figure 1E). Finally, we generated a complete whole-brain cell type atlas in CCFv3 space at 10 µm isotropic resolution (Figure 1F). This study integrates high-resolution dMRI to CCFv3 space, providing high spatial resolution water diffusivity and brain microstructural information that can be further compared across different modalities. This study also demonstrates the usage of dMRI in predicting different taxonomy levels of neuronal cell types in the adult mouse brain. The resulting complete whole-brain cell type atlas may serve as a detailed visualization of brain structures and functions, enabling insights into the organization of neural circuits, developmental processes, and the underlying mechanisms of neurological diseases.

**Figure 1.**
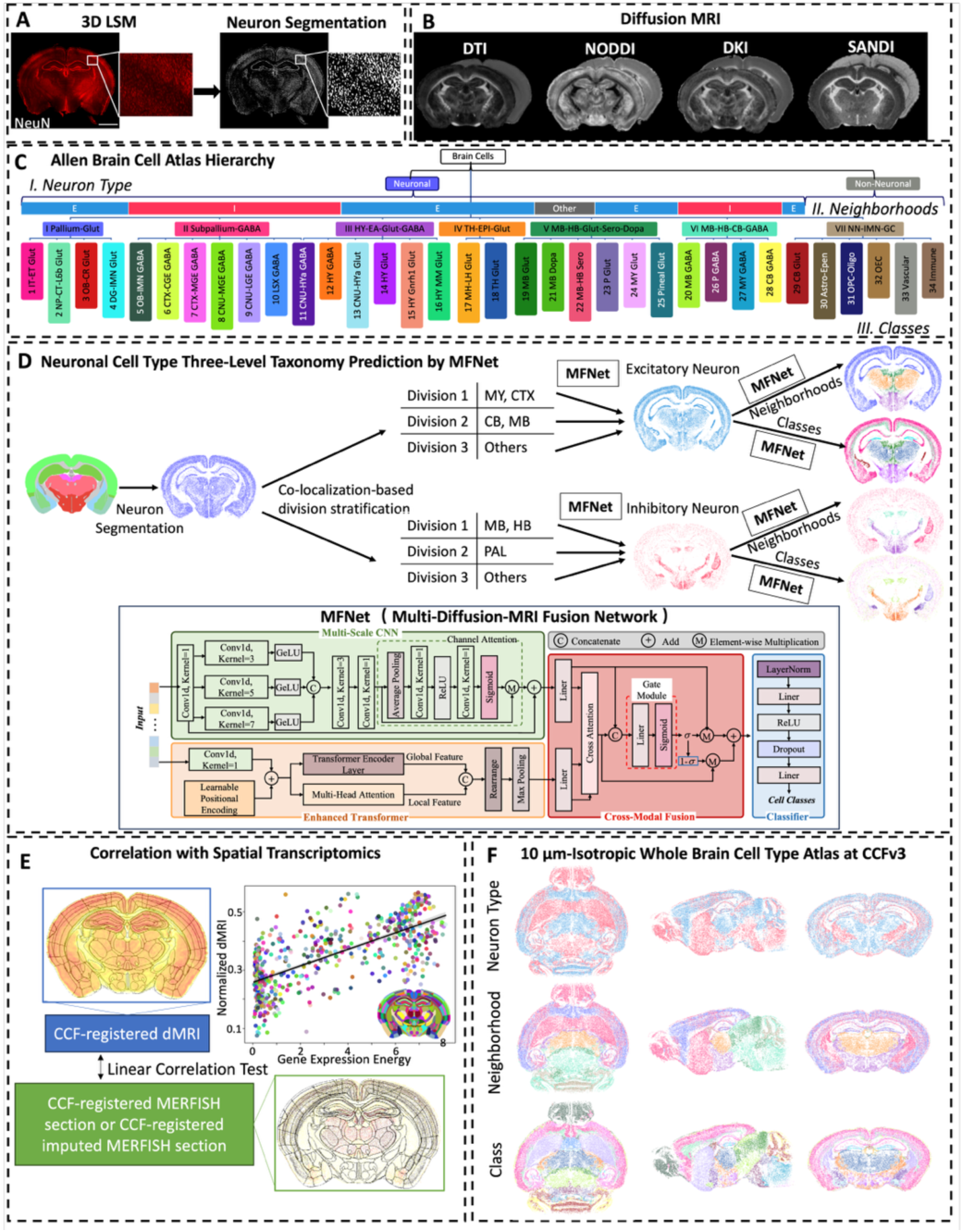
Overview of this study. (A) Neuron segmentation from a high-resolution 3D NeuN image. Scale bar = 2µm. (B) High-resolution diffusion MRI images. Left to right: dMRI metrics from DTI, NODDI, DKI, and SANDI models. (C) The transcriptomic taxonomy tree of Allen Brain Cell (ABC) Atlas with level I: Neuron Type, level II: Neighborhoods, and level III: Classes. (D) Cell type classification model by MFNet. Top: Overview of the classification process. The slice is color-coded by the three hierarchical levels. Bottom: MFNet Architecture. (E) Correlation test of dMRI images (in blue box) with spatial transcriptomics (in green box), both registered in CCF space (CCF annotation boundary overlaid). (F) The generated complete whole-brain cell type atlas at CCFv3 space. The cell type atlas is color-coded by neuron type (top), cell neighborhood (center), and cell class (bottom), shown on example axial (left), sagittal (middle), and coronal (right) views.

## 2. Results

### 2.1. MRI and LSM integrated into the CCF space

A multi-modality brain map (Table S1), including dMRI maps with fourteen different image contrasts (Figure 2B and C) and 3D LSM images (Figure 2A), has been integrated into CCF space. The alignment accuracy that was measured by the Dice similarity coefficient of each pair of spatially matched coronal slice (n = 1320) between dMRI and CCF is 0.96 ± 0.06, and between 3D LSM and CCF is 0.92 ± 0.09. The robust alignment enabled accurate downstream analyses and yielded a multimodal atlas in a common coordinate framework.

**Figure 2.**
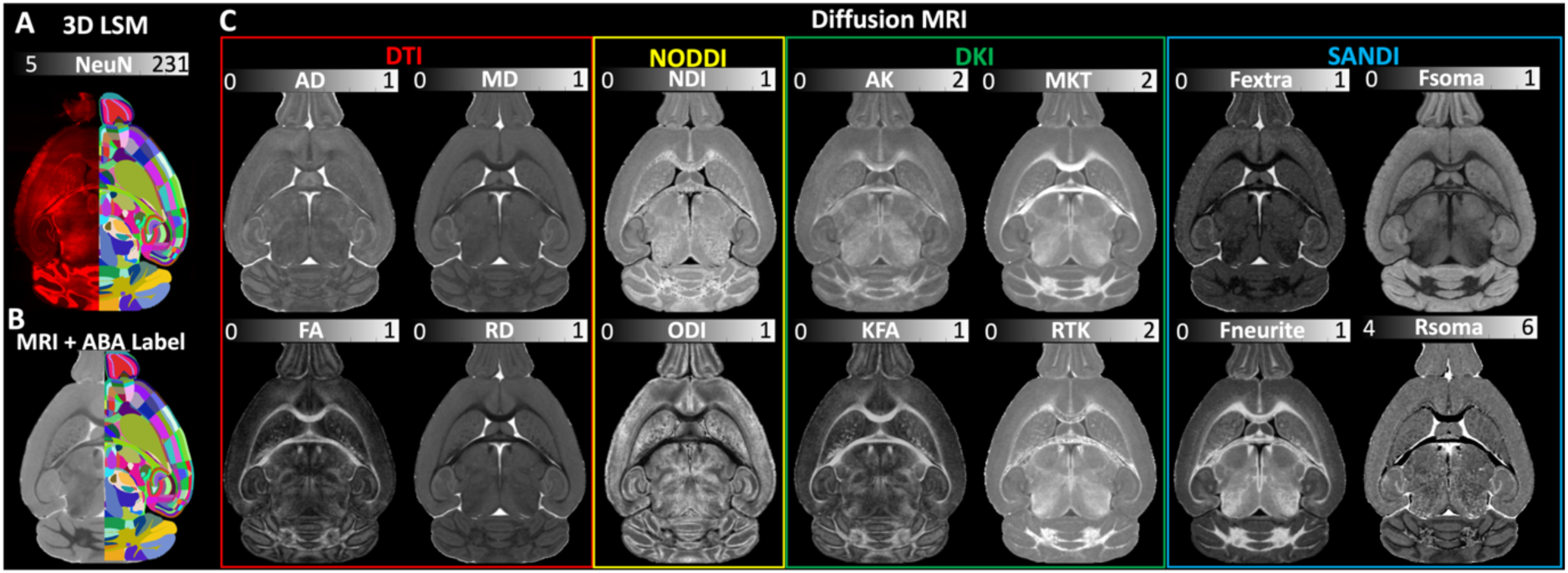
CCF space integrated dMRI and 3D LSM. (A) NeuN image overlaid with ABA label. (B) B_0_ image overlaid with ABA label. (C) dMRI maps, including metrics from DTI (in red box), NODDI (in yellow box), DKI (in green box), and SANDI (in blue box) models. See Figure S1 for the dMRI maps in native space.

### 2.2. Whole-brain cell type prediction evaluation

Using CCF-integrated dMRI, we predicted the spatial distribution of neuronal cell types across the whole mouse brain. MFNet outperformed baseline deep learning models, a multi-layer perceptron (MLP) (Table S2). Division-specific training further improved performance compared to a unified model (Figure S2). Within MFNet, the multi-scale CNN provided the primary predictive signal, the Transformer supplied complementary contextual information, and cross-modal fusion was essential for achieving these gains, particularly in cell type classification. Training and validation loss curves, together with performance metrics, confirmed that MFNet is both effective and generalizable (Figure S3).

Figure 3 shows the prediction results in the hippocampus (HPF) and isocortex regions, and the whole brain. In the HPF, the weighted average prediction accuracy is 88.8±6.3% for the excitatory neuron type (Figure 3A, left), 94.8±0.01% for the inhibitory neuron type (Figure 3B, left). In the isocortex, the weighted average prediction accuracy is 87.7±0.9% for the excitatory neuron type (Figure 3A, right), 94.8±0.2% for the inhibitory neuron type (Figure 3B, right). Across the whole brain, MFNet predicted cell neighborhoods (Figure 3C) better than cell classes (Figure 3D); the weighted average accuracy is 97.8±2.9% / 98.6±2.0% for E/I neighborhood predictions (Figure 3C) and 88.6±3.9% / 82.6±10.9% for E/I class predictions (Figure 3D). Overall, there is no significant difference between the performance of MFNet in excitatory and in inhibitory neuron types (Figure 3E).

**Figure 3.**
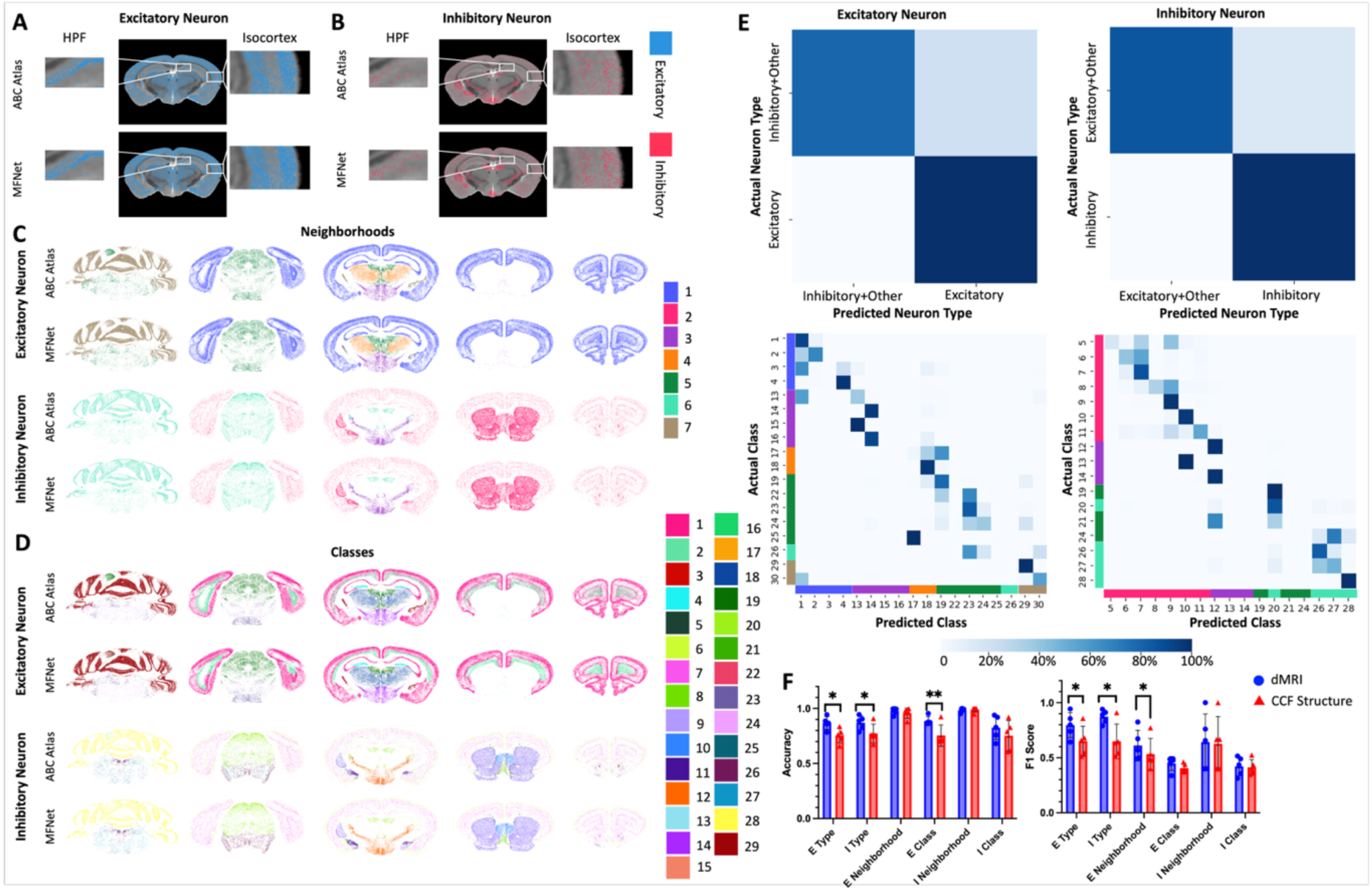
Neuronal cell type prediction validation. (A, B) Prediction results on excitatory neuron type (A) and inhibitory neuron type (B) in whole brain (middle), HPF (zoomed left), and isocortex (zoomed right) regions. Top: ABC Atlas; bottom: MFNet predicted. (C, D) Prediction results on neuronal cell neighborhoods (C) and classes (D) in the mouse whole brain from each one of the testing slices. From top to bottom: excitatory neurons from ABC Atlas and MFNet predicted, inhibitory neurons from ABC Atlas and MFNet predicted. (E) Confusion matrices between MFNet predicted excitatory neuron type and actual (top left), between MFNet predicted excitatory cell classes and actual (bottom left), between MFNet predicted inhibitory neuron type and actual (top right), and between MFNet predicted inhibitory cell classes and actual (bottom right). Bar graphs on the right and bottom of the confusion matrix indicate the corresponding cell neighborhoods. (F) Bar graph comparing the weighted average prediction accuracy (left) and F1 score (right) in MFNet-dMRI (in blue bars) and MFNet-CCF structure (in red bars). Error bars represent the standard deviation among the testing slices. n = 5. * FDR-corrected p-values < 0.05, ** FDR-corrected p-values < 0.01. The results shown in this figure were obtained using MRI data from the high b-value protocol. See results using the high angular and spatial resolution protocols in Figure S5 and S6.

To investigate the importance of dMRI over anatomical locations in predicting cell types, we compared the performance of MFNet using dMRI or CCF structures (*i.e.,* baseline feature, as the ABC Atlas cell types are defined based on their location in the CCF broad region). From MFNet-CCF structure (Figure 3F, red bars) to MFNet-dMRI (Figure 3F, blue bars), the performance has increased substantially in neuron type predictions (+ 15.1% / 12.8% on accuracy and + 22.4% / 34.7% on F1 score) and excitatory cell class predictions (+ 17.6% on accuracy). The most significant increases occurred in the ventricular systems (VS), pallidum (PAL), fiber tracts, and unassigned regions. In contrast, they performed similarly in E/I neighborhood and inhibitory cell class predictions. We also compared the performance of MFNet with and without the cell coordinates and found that excluding the coordinates caused 12.4% / 24.3% decreasing on E/I class prediction only (Figure S4). Furthermore, we evaluated the robustness of MFNet for image acquisition setting dependency by training with high angular and spatial resolution protocol. MFNet showed a consistently high degree of prediction accuracy (Figure S5 and S6).

### 2.3. Diffusion MRI can distinguish cell type

Diffusion MRI contained sufficient information to classify neuronal cell types. UMAP projections (Figure 4A) revealed distinct spatial clustering across the whole brain (top), isocortex (center), and HPF (bottom), regarding neuron type (column 2), excitatory cell neighborhood (column 3) and class (column 4), inhibitory cell neighborhood (column 5) and class (column 6). This demonstrates that diffusion MRI encodes key microstructural information for cell type discrimination. Heatmaps of dMRI metrics (Figure 4B) further showed cell–type–specific patterns, with SANDI-derived metrics exhibiting the strongest associations with cell type distributions. Feature importance analysis identified Fsoma and Fneurite as the top predictors (Figure S7), and both metrics displayed clear distinctions across neuron types, neighborhoods, and classes (Figure 4C).

**Figure 4.**
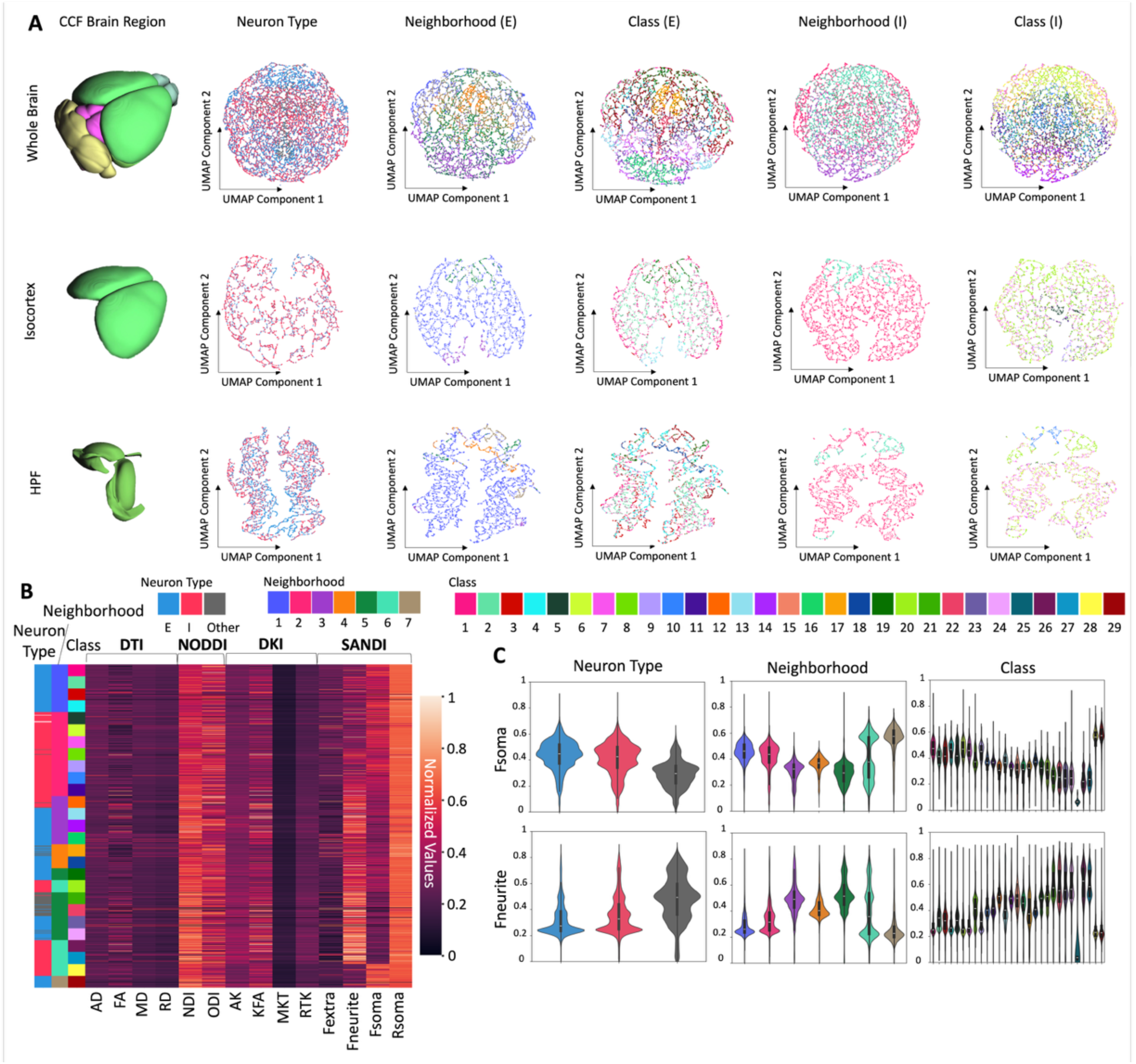
(A) U-MAP of cell neighborhoods in the whole brain (top), isocortex (center), and HPF (bottom), color-coded by neuron type (column 2), excitatory cell neighborhood (column 3) and class (column 4), inhibitory cell neighborhood (column 5) and class (column 6). (B) Heatmap showing the normalized dMRI values distributed in different cell types. Bar graphs on the left show the neuron type (left), cell neighborhood (middle), and class (right). (C) Violin plots showing the distribution of Fsoma (top) and Fneurite (bottom) in different neuron types (left), cell neighborhood (middle), and class (right).

### 2.4. Diffusion MRI is correlated to the spatial transcriptomics of neuron-type marker genes

Diffusion MRI predicts cell types effectively because its metrics correlate with spatial transcriptomic profiles of cell type marker genes. Figure 5 shows the correlation between dMRI metrics and spatial transcriptomics. Specifically, the expression profiles of the neuron type marker genes *Slc17a6*, *Slc17a7*, *Slc32a1*, *Gad1*, *Gad2*, and *Aldh1a1* showed similar spatial patterns to dMRI metrics in the heatmap (Figure 5A). Particularly for the gene *Slc17a7* (excitatory neuron type marker gene) and *Aldh1a1* (inhibitory neuron type marker gene), their high Pearson’s correlation coefficients (PCC) with most of the dMRI metrics demonstrated such strong correlations (Figure 5B, right). Among the *Slc17a7*-dMRI and *Aldh1a1*-dMRI correlations, *Slc17a7/Aldh1a1*-Fsoma and *Slc17a7/Aldh1a1*-Fneurite were the most significant ones (Figure 5C). For example, in the isocortex (black arrows) and HPF (blue arrows) regions (Figure 5D), *Slc17a7* is highly expressed, whereas *Aldh1a1* shows lower expression. Correspondingly, Fsoma values are high and Fneurite values are low in these regions. Across all genes, FA, NDI, AK, KFA, MKT, RTK, and Fneurite were predominantly negatively correlated, whereas AD, MD, RD, ODI, Fextra, Fsoma, and Rsoma showed predominantly positive correlations. (Figure 5B, left).

**Figure 5.**
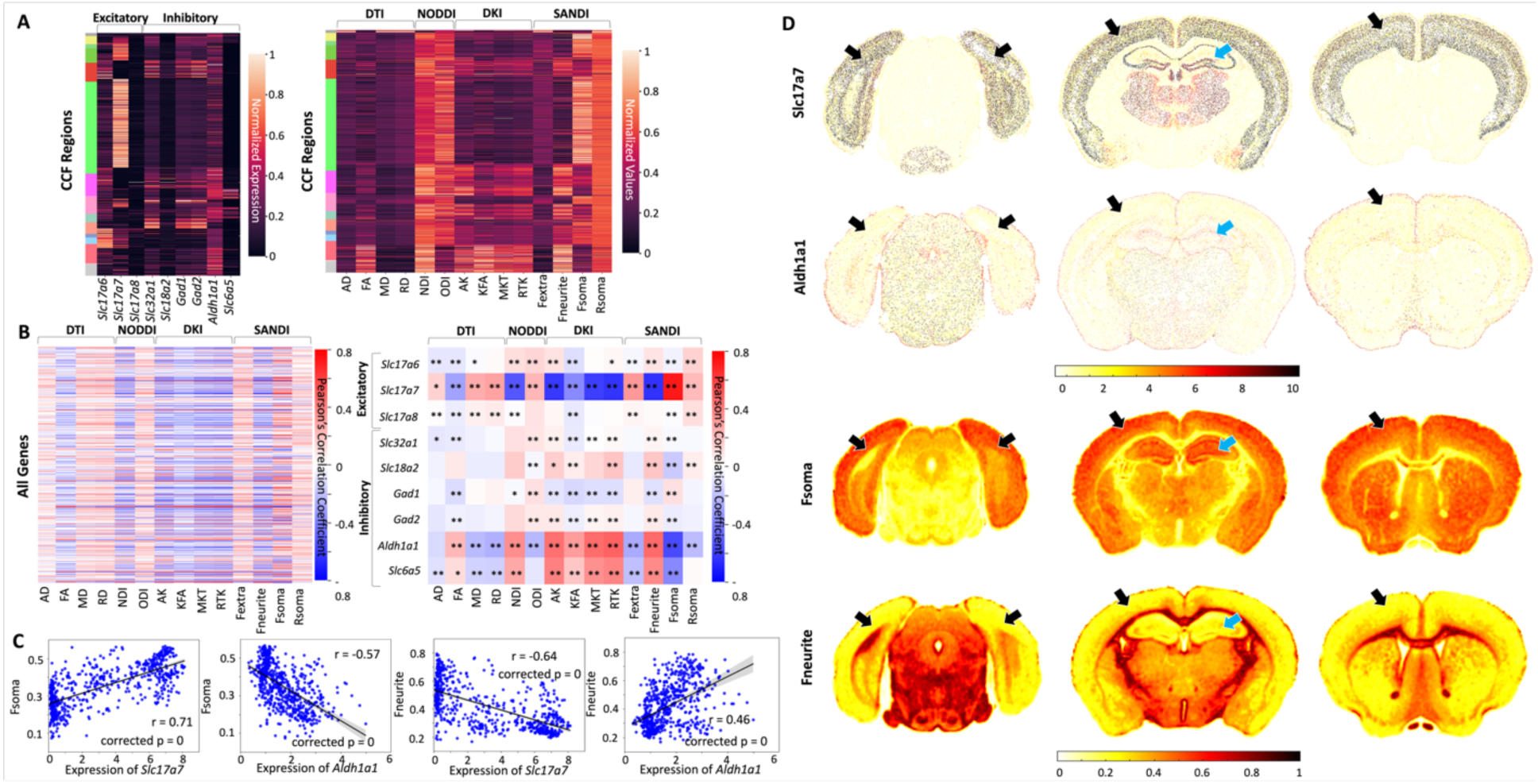
dMRI metrics correlation with spatial transcriptomics of MERFISH gene panels. (A) Heatmap comparing the normalized expression energy of neuron type marker genes (left) and the normalized dMRI values (right) distributed in CCF regions. The row represents one CCF annotation region. Bar graphs on the left show the broad CCF regions. (B) Heatmap of PCC between dMRI parameters and the spatial transcriptomics of all MERFISH genes (left) and neuron type marker genes (right). Each row represents one gene. * Spatial autocorrelation (SA)-corrected p-values < 0.05, ** SA-corrected p-values < 0.01. (C) Linear regression plots of four independent variable-dependent variable pairs (left to right): Slc17a7 (x axis)-Fsoma (y axis), Aldh1a1 (x axis)-Fsoma (y axis), Slc17a7 (x axis)-Fneurite (y axis), Aldh1a1 (x axis)-Fneurite (y axis). n = 686. The PCC r values, SA-corrected p-values, and 95% confidence intervals (in grey shadows) were shown in each plot. (D) Representative slices of the spatial transcriptomics of neuron type marker gene Slc17a7 (row 1) and Aldh1a1 (row 2), as well as dMRI Fsoma (row 3) and Fneurite (row 4) maps. Black arrows and blue arrows indicate the isocortex and HPF, respectively.

### 2.5. MFNet Atlas covers the whole brain volume and is more complete compared to the ABC Atlas

Using MFNet, we generated a whole-brain mouse cell type atlas, termed the **MFNet Atlas** (Figure 6), in CCFv3 space at 10 µm isotropic resolution, with annotations across all three taxonomic levels. Compared to the ABC Atlas, the MFNet Atlas exhibits comparable anatomical and spatial detail (Figure 6A), with average similarity scores across 35 spatially matched slices of 0.99 ± 0.01 / 0.98 ± 0.02 for E/I neuron type, 0.98 ± 0.02 / 0.98 ± 0.02 for E/I neighborhood, and 0.99 ± 0.01 / 0.93 ± 0.05 for E/I class; observed discrepancies (arrows in Figure 6A) primarily reflect differences in neuron segmentation (Figure S8A).

The MFNet Atlas also offers several key advantages over the ABC Atlas. First, 34% of slices in the ABC Atlas were damaged, resulting in missing cell type information (arrows in Figure 6B and Figure S8B), whereas MFNet recovers these regions to provide a complete atlas. Second, MFNet achieves higher accuracy than the interpolated ABC Atlas; for example, inhibitory neuron distributions in regions highlighted by arrows (Figure 6C) show greater concordance with Gad1 and Gad2 (two common inhibitory neuron marker genes) *in situ* hybridization (ISH) ground truth. Third, MFNet provides whole-brain coverage, including 1320 coronal, 1140 sagittal, and 800 axial views, compared to 53 coronal views in the ABC Atlas. MFNet generates a continuous 3D volume, whereas the ABC Atlas is limited to discrete 2D slices (Figure 6D). In summary, the MFNet Atlas delivers more complete and accurate cell type information than the ABC Atlas, establishing a high-resolution resource for investigating the brain at the whole-volume level.

**Figure 6.**
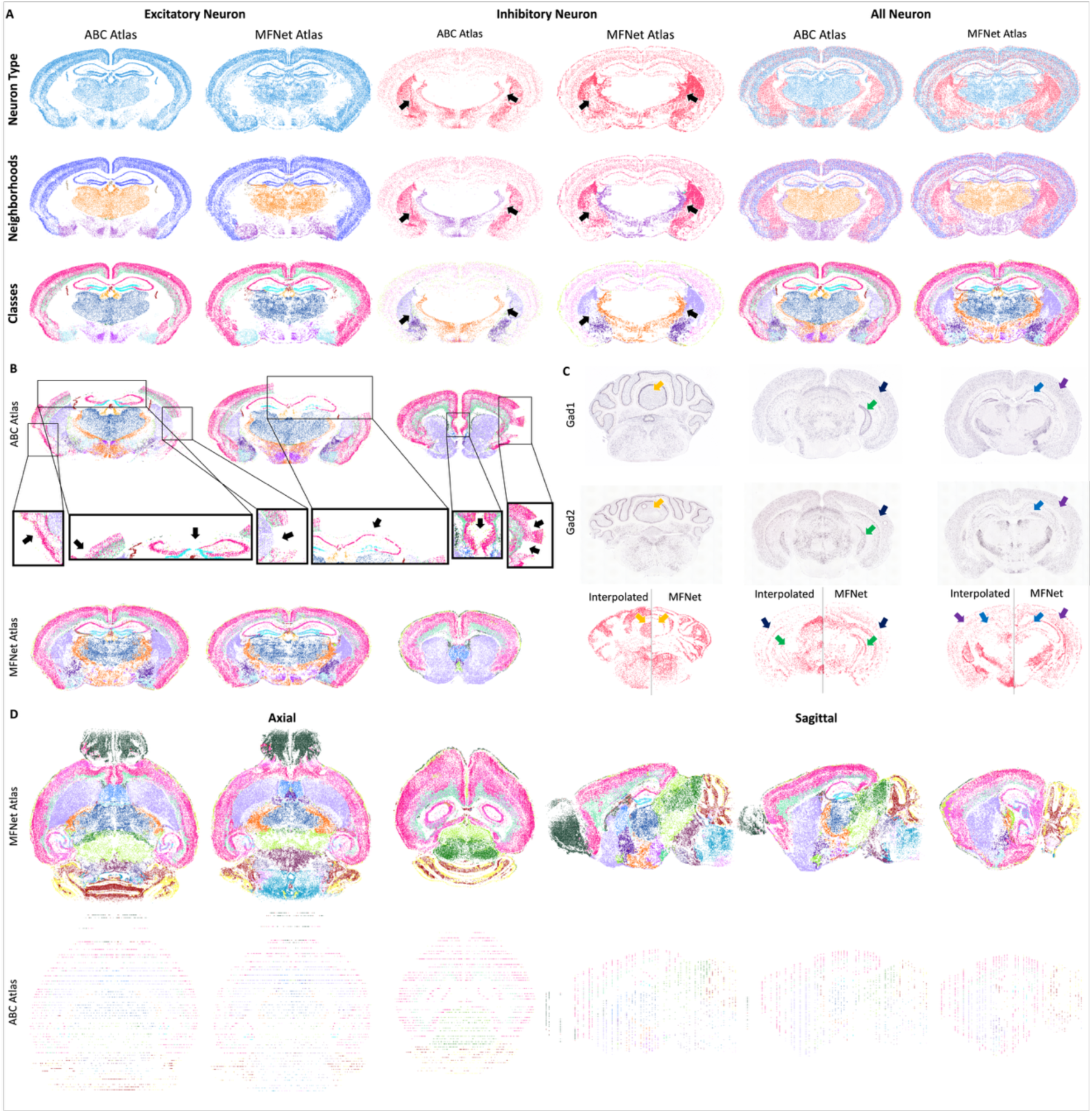
MFNet Atlas, a complete, whole mouse brain cell type atlas at CCFv3 space with 10 μm isotropic resolution. (A) Comparisons between one representative slice from the MFNet Atlas (right) and the corresponding ABC Atlas (left). The two atlases are color-coded by neuron type (columns 1∼3), cell neighborhoods (columns 4∼6), and classes (columns 7∼9), with excitatory neurons (top) and inhibitory neurons (center) shown separately, as well as all neurons (bottom). Arrows indicate the major mismatched areas. (B) Comparisons between three representative slices from the MFNet Atlas (bottom) and the corresponding ABC Atlas (top) that were damaged. The two atlases are color-coded by cell class of all neurons. Arrows in the enlarged regions indicate the damaged areas. See comparisons on more slices in Figure S6. (C) Comparisons between three representative slices from MFNet Atlas (right) and corresponding interpolated ABC Atlas (left) with reference Gad1(top) and Gad2 (center) ISH images (Allen Mouse Brain Atlas, mouse.brain-map.org). The two atlases are color-coded by neuron type of inhibitory neuron. Arrows indicate the spatially matched areas, including cerebellar cortex, isocortex, and hippocampus. (D) Representative slices from axial (left three) and sagittal (right three) views from the MFNet Atlas (top) and corresponding ABC Atlas (bottom) are shown here. Under each view, the atlas is color-coded by cell class of all neurons.

## 3. Discussion

Understanding the structure and function of the brain remains one of the most challenging topics in contemporary neuroscience. Significant efforts have been devoted to refining high-resolution imaging techniques for more detailed anatomical insights, ^24,25^ while simultaneously other efforts have focused on creating cellular taxonomies to elucidate underlying mechanisms.^26,27^ Integrating imaging techniques with taxonomy findings will significantly enhance the understanding of the brain at different scales.^28^ In this study, we integrated MRI images and LSM into the Allen Mouse Brain Common Coordinate Framework, providing an atlas with both structural and functional information for the whole mouse brain. Compared to the previous studies ^29,30^ that were solely on the basic DTI model, this integrated atlas incorporates both DTI and advanced diffusion models, enabling a more precise characterization of tissue microstructure. Furthermore, it provides the neuroimaging community with a high-resolution multi-parameter MRI template in the CCFv3 space with detailed cell types across the whole brain.

The predictive power of dMRI for cell types arises from its sensitivity to anatomical and functional features. dMRI detects microstructural changes, including fiber orientations in white matter,^31^ cortical soma size and density, and fine cortical and olfactory bulb structures.^12^ High-resolution dMRI also reveals neuroinflammation,^32^ neurodegeneration,^16^ glial activation,^33^ and cellular-level tissue features such as cell morphology, membrane integrity, and local cell density.^34,35^ dMRI differentiates intracellular and extracellular space as intracellular molecules generally diffuse much more slowly than extracellular molecules,^36^ allowing for the showing up of high cell density areas with high intensity. Beyond microstructure, diffusion metrics such as NDI and ODI correlate with gene expression profiles, linking MRI signals to molecular pathways.^37,38^ Spatial imaging-transcriptomics further confirms that regional diffusion markers reflect underlying gene expression patterns, highlighting dMRI as a non-invasive probe of cellular and genetic features.^21^ Our previous study also demonstrated that dMRI metrics are associated with the genes involved in brain development.^39^

MRI outperforms CCF-based anatomical labels in distinguishing neuronal cell types. While CCF regions exhibit regional specificity and contain distinct neuronal populations, many cell types—including glutamatergic and GABAergic neurons—colocalize within the same regions, limiting the discriminatory power of anatomical labels alone. Differentiating these colocalized cell types, therefore, requires higher-order structural and functional information, which MRI provides at high resolution. In contrast, high-resolution 3D LSM primarily captures spatial distributions of neurons.^40,41^ To leverage the strengths of both modalities, we assigned complementary roles: cell segmentation was performed using LSM data, while cell type prediction relied on MRI data. Integrating these results enabled the generation of a high-resolution neuronal cell type atlas spanning the entire mouse brain.

Notably, SANDI parameters—particularly Fsoma and Fneurite—ranked as the top two features among all dMRI metrics for predicting cell types. This is consistent with the design of the SANDI model, a biophysical framework specifically developed to estimate cellular and neurite densities,^18^ rendering it highly sensitive to cellular and molecular differences between neuronal types. More specifically, Fsoma describes the fraction of soma that produces neurotransmitters, while Fneurite would signal the presence of a prominent apical dendrite on cortical pyramidal neurons.^18^ Given that neuronal cell types are defined by their neurotransmitter types in the ABC Atlas, the high predictive power of Fsoma and Fneurite can also be attributed to their strong correlation with the spatial transcriptomics of neurotransmitter marker genes. *Slc17a7* encodes Vesicular Glutamate Transporter 1 (VGLUT1), a key protein for Glut neurotransmission.^42,43^ *Aldh1a1* is crucial both for GABA metabolism and for enabling dopaminergic neurons to co-release GABA.^44,45^ As shown in Figure 5, both Fsoma and Fneurite correlate strongly with the spatial expression of these marker genes, supporting their role in distinguishing neuronal cell types.

Leveraging these insights, we generated a 3D, whole-brain MFNet Atlas with 10 µm isotropic resolution in CCFv3 space. Existing mouse brain cell type atlases offer high in-plane resolution but limited axial resolution (200 µm),^3,6^ resulting in incomplete coverage across many brain regions. Leveraging the high isotropic resolution of diffusion MRI, we generated a whole-brain, MRI-guided cell type atlas with 10 µm axial resolution, fully consistent with CCFv3.^46^ The MFNet Atlas also provides micrometer-scale in-plane resolution (10 µm), comparable to the average size of a mouse brain cell.^47^ A key advantage of MFNet is its ability to accurately annotate cell types in regions where the ABC Atlas is incomplete, including damaged or intervening slices. Beyond completeness, the MFNet Atlas enables studies of brain development, evolution, function, and pathology,^3,48,49^ facilitating detailed investigation of molecular and anatomical architecture, characterization of morphological and physiological properties, and precise spatial mapping of distinct neuronal populations.

Diffusion MRI–based cell type prediction has broad research and clinical utility. Traditionally, characterizing cell types requires labor-intensive approaches, including transcriptomic profiling, molecular identification, developmental analyses, and constructing conceptual frameworks for cell type definition.^4^ In contrast, deep learning–based prediction models can infer cell types without extensive domain expertise, and high-resolution MRI enables cellular-level microstructural information to be captured. Given that diffusion MRI is already widely applied in studies of brain disorders,^16,50^ integrating cell type prediction into existing imaging workflows could provide additional cellular-level insights, enhancing both clinical and translational research. The MFNet framework can also be used to quantify the excitatory/inhibitory (E/I) ratio, which is critical for maintaining proper brain function and network stability.^51^ Dysregulation of this balance has been implicated in neurodevelopmental disorders^52^ and is also an early hallmark of AD progression in mouse models.^53^ Therefore using MFNet prediction framework may assist early detection and treatment assessment of neurological disease.

Several limitations remain. First, the number of mice was limited due to lengthy image acquisition times. Second, the prediction model currently cannot be applied to non-neuronal cells, which are widely distributed, highly colocalized, and less region-specific, leading to similar MRI values that are difficult to differentiate. Moreover, the raw MRI resolution is substantially lower than that of MERFISH,^54^ resulting in near-identical MRI values among neighboring cells and limiting classification performance. Third, the accuracy of MFNet partly depends on prior knowledge of the cellular composition characteristic of the genotype. For example, the model infers cell-type proportions from reference labels, which may be less reliable in slices that do not match the ABC Atlas data or that originate from different genotypes. In addition, the brain-division stratification step relies on predefined degrees of cell-type co-localization, information that may be unavailable or inaccurate for some genotypes. Fourth, although current dMRI achieves microscopic-level resolution, it remains substantially coarser than the 10 µm atlas. Further increasing spatial resolution could enhance the accuracy of the predicted cell-type atlas. Last, neuron density in the MFNet Atlas is higher than in the ABC Atlas, primarily because MFNet utilizes neuronal segmentation directly from 3D NeuN images, whereas the ABC Atlas has excluded low-quality cells to improve sequencing accuracy.^3^ Future improvements could include enhancing the spatial resolution of dMRI, refining models for non-neuronal cell types, applying unsupervised or transfer learning approaches to blank slices, employing spatially independent models, and integrating sparse cell distributions into the MFNet Atlas.

## 4. Conclusion

In this study, we integrated high-resolution diffusion MRI and three-dimensional light-sheet microscopy within the Allen Mouse Brain Common Coordinate Framework to generate a comprehensive neuronal cell type atlas. A deep learning–based network accurately predicted neuronal cell types across the entire mouse brain, demonstrating that dMRI parameters capture key cellular properties and strongly correlate with the spatial distribution of neurons. We produced a 3D whole-brain atlas with high in-plane and through-plane resolution. Collectively, these results highlight the power of advanced imaging techniques for mapping brain structure and function and provide a high-resolution resource for understanding the brain at cellular and molecular scales.

## 5. Methods and Materials

### 5.1. Animal study

Six adult C57BL/6J (B6, wild type) male mice (Jackson Laboratory, Bar Harbor, ME) were selected for MR imaging. All animal experiments were conducted in compliance with the Duke University Institutional Animal Care and Use Committee (Approval code A226-17-09). The mice were euthanized and perfusion-fixed with a 1:10 mixture of ProHance-buffered (Bracco Diagnostics, Princeton, NJ) formalin. The specimens were immersed in buffered formalin for 24 hours and then transferred to a 1:200 solution of ProHance/saline to shorten the T1 (to approximately 100 ms) and reduce scan time. *Ex vivo* MRI was chosen over an *in vivo* study to achieve higher anatomic resolution and better tissue contrast, which improves the detection of microstructural changes^55,56^ and to maintain consistency with *ex vivo* spatial transcriptomics data.

### 5.2 Allen Mouse Brain Atlas processing

Allen Mouse Brain Atlas (AMBA)’s serial two-photon tomography (STP) image templates and 3D annotation labels in Allen Mouse Brain Common Coordinate Framework with 10 μm isotropic resolution (CCFv3) and resampled CCF (rCCF) mapped coordinates with 10 μm in-plane and 200 μm through-plane resolution were downloaded from the Allen Brain Cell (ABC) Atlas website (https://portal.brain-map.org/atlases-and-data/bkp/abc-atlas) and converted to NIFTI format.^46^

To keep the pixel size consistent between the CCFv3 ( x × y × z = 800 × 1140 × 1320) and rCCF ( x × y × z = 1100 × 1100 × 76) space, we firstly up-sampled the z-axis of STP image at rCCF space from 76 to 1100, and then we registered the STP image from CCFv3 to up-sampled rCCF space using Advanced Normalization Tools (ANTs)’ affine image registration. The transformation matrix was applied to CCFv3’s 3D annotation labels. The CCFv3 space in this study all refers to the registered one.

### 5.3. MRI processing

#### 5.3.1. MRI acquisition

All the MRI experiments were performed on a 9.4T Oxford 8.9-cm vertical bore magnet (Oxford Instruments, Abingdon, United Kingdom), and the maximum gradient amplitude on each axis was 2000 mT/m. A modified three-dimensional (3D) diffusion-weighted spin-echo pulse sequence with k-space undersampling was used in this study. The readout dimension was fully sampled, and both phase encoding dimensions were undersampled using a sparsifying approach, described in detail previously.^24^ An acceleration factor (AF) of 4.0 was used in this study, where 1.0 stands for the fully sampled data. For multi-shell dMRI acquisition, the diffusion gradient directions were determined using the method proposed by Koay et al.^57^ to ensure uniformity within each shell while maintaining uniformity across all shells.

The dMRI under sampled datasets were acquired with three protocols: 1) high-b value protocol (n = 1): 16 shells b value = 1000, 2000, 3000, 4000, 5000, 6000, 7000, 8000, 9000, 10000, 11000, 12000, 13000, 14000, 15000, 16000 s/mm^2^ at 45 μm isotropic resolution, gradient separation (GS) time of 10 ms, gradient duration (GD) time of 4.8 ms, and the total scan time was approximately 72 hours. 2) high angular resolution protocol (n = 1): 8 shells b value = 1000, 2000, 3000, 4000, 5000, 6000, 7000, 8000 s/mm^2^ at 50 μm isotropic resolution, GS time of 7.7 ms, GD time of 4.8 ms, and the total scan time was approximately 148 hours. 3) high spatial resolution protocol (n = 4): 4 shells b value = 1000, 4000, 6000, 8000 s/mm^2^ at 32 μm isotropic resolution, GS time of 7.4 ms, GD time of 4.8 ms, and the total scan time was approximately 48 hours. 64 diffusion-weighted images (DWIs) were acquired in each shell. The repetition time (TR) was 100 ms, the echo time (TE) was 16.65 ms, the bandwidth (BW) was 156 kHz, and the maximum gradient amplitude was about 140 mT/m.

#### 5.3.2. Compressed sensing

The under-sampled k-space data reconstruction has been described in previous studies^24,58^ by minimizing the following function:

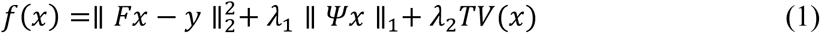

Where 𝑥 is the image and 𝑦 is its corresponding k-space, *F* is the FFT, 𝛹 is the sparse transform, 𝜆_1_ and 𝜆_2_ are regularization parameters, and 𝑇𝑉 is the total variation. In this study, 𝜆_1_ equals 0.006 for the sparse solution and 𝜆_2_ equals 0.0012 for the data consistency with the Daubechies wavelet transform performed^58^. An acceleration factor of 4.0 means the acquisition time is 1/4_th_ of the time required for a fully sampled dataset. The total scan time of the multi-shell dMRI datasets was approximately 48 hours; the scan time for the fully sampled dataset would be 192 (48 × 4) hours. The reconstruction was implemented on a Dell Cluster, where an initial Fourier transform along the readout axis yielded multiple 2D images from each acquisition that could be spread across the cluster for parallel reconstruction.

#### 5.3.3. DTI metrics extraction

Diffusion Tensor Imaging (DTI) reconstruction method was performed for reconstruction using DSI Studio ^59^. From data with b values of 1000 to 4000 s/mm^2^, the diffusion tensor was calculated. The scalar indices include fractional anisotropy (FA), mean diffusivity (MD), axial diffusivity (AD), and radial diffusivity (RD).

#### 5.3.4. NODDI metrics extraction

The NODDI toolbox for Matlab was used to fit the diffusion MRI data with b values of 1000 to 8000 s/mm^2^.^60^ For the *ex vivo* NODDI model, a relatively small dot-compartment (the diffusion is restricted in all directions) was included for the fitting.^61^ The parameters derived from the NODDI model were the orientation dispersion index (ODI) and neurite density index (NDI).

#### 5.3.5. DKI metrics extraction

Diffusion Imaging in Python (Dipy, version 1.10.0) was used to compute DKI metrics from data with b values of 1000 to 8000 s/mm^2^. Self-Supervised Denoising via Statistical Independence (Patch2Self)^62^ was used before fitting to the DKI model for image denoising. The DKI model has been set up to use constraints with the option fit_method = ‘CLS’ (for ordinary least squares) and cvxpy_solver = ‘MOSEK’. The DKI parameters generated were axial kurtosis (AK), kurtosis fractional anisotropy (KFA), the mean kurtosis tensor (MKT), and radial tensor kurtosis (RTK).

#### 5.3.6. SANDI metrics extraction

SANDI Matlab Toolbox^18^ was used to calculate SANDI metrics from data with all b values. Denoising using “dwidenoise” from MRTrix3 was used to save the estimated noisemap. The SANDI metrics recorded for further analysis were soma fraction (Fsoma), neurite fraction (Fneurite), radius of soma (Rsoma), and ratio of neurite (Rneurite).

#### 5.3.7. MRI registration

The diffusion MRI images were registered to the CCFv3 space and the rCCF space using ANTs’ affine and syn image registration, which has been described in a previous study in detail.^63^

### 5.4. Histology image processing

#### 5.4.1. Three-Dimensional LSM image and in situ hybridization image acquisition

The 3D NeuN staining (Fox3/NeuN, mouse monoclonal, Cat# MCA-1B7, RRID# AB_2572267), images (n=3) and 3D GAD67 staining (GAD67 Monoclonal Antibody (CL2914), Cat# MA5-31377, RRID# AB_2787014) image (n=1) for adult male B6 mice were downloaded from the Duke Mouse Brain Atlas (DMBA, https://civmimagespace.civm.duhs.duke.edu/)^41^ and converted to NIFTI format. The Gad1 and Gad2 *in situ* hybridization (ISH) staining images for adult male B6 mice were downloaded from Allen Mouse Brain Atlas (https://mouse.brain-map.org/. Experiment # 479 and # 79591669, respectively)Click or tap here to enter text..

#### 5.4.2. Neuron cell segmentation and pre-processing

Neuron cells were segmented from 3D NeuN images at their native space. Labkit ^65^ automatic segmentation was used for each image slice (Figure 1A). The segmentation result images were saved as .tiff format, and were pre-processed by firstly averaging per genotype group and secondly binarizing using a locally adaptive threshold.

#### 5.4.3. Image registration

The 3D NeuN images were registered to CCFv3 space using ANTs’ affine and syn image registration. The registration was refined manually guided by anatomical landmarks such as hippocampal subfields and cortical layers. The transformation matrices were applied to their corresponding pre-processed segmentation result images.

### 5.5. Mouse whole-brain transcriptomic cell type prediction model

#### 5.5.1. Transcriptomic cell type atlas data

The mouse whole-brain transcriptomic cell type atlas consisted of whole adult mouse brain MERFISH spatial transcriptomics and taxonomy of cell types dataset, which were exported from Allen Brain Cell Atlas (ABC Atlas, https://portal.brain-map.org/atlases-and-data/bkp/abc-atlas). The MERFISH spatial transcriptomics dataset included the coordinate information of each cell centroid from the whole mouse brain. Such coordinates were provided in rCCF space. The taxonomy dataset integrated several whole-brain single-cell RNA-sequencing (scRNA-seq) datasets and contains cell type clusters in a hierarchical structure. Four levels of such structure were selected in this study, including (from top to bottom): neuronal/non-neuronal cells, 3 neuron type (excitatory (E): Glut neurons /inhibitory (I): GABA, GABA-Glyc, and Glut-GABA neurons/others: Chol, Dopa, Hist, Nora, and Sero neurons), 7 cell neighborhoods, and 34 cell classes (Figure 1C). All the selected levels followed the same hierarchy as the ABC Atlas. Click or tap here to enter text. Non-neuronal cell types and other neuron types were deleted from the data as they are out of the scope of the current topic.

#### 5.5.2. Data pre-processing for model training Training/testing dataset split

A total of 53 MERFISH section slices (MERFISH-C57BL6J-638850.05∼ MERFISH-C57BL6J-638850.67 Reconstructed Coordinates) were used, covering the whole anterior-to-posterior brain volume. We selected one section slice from each fifth of the brain volume as a testing section slice (MERFISH-C57BL6J-638850.14, MERFISH-C57BL6J-638850.26, MERFISH-C57BL6J-638850.36, MERFISH-C57BL6J-638850.51, and MERFISH-C57BL6J-638850.58), with the remaining section slices used as training data. This will ensure that the testing data is representative of the entire dataset and independent of the training data.

##### Brain division stratification according to the co-localization level of neuron types

Heterogeneous colocalization levels were observed in the spatial distribution of E/I neuron types across particular brain regions.^3^ In alignment with this observation, we developed a division-specific neuron type prediction model (Figure 1D) to separately train the whole brain into three brain divisions for each E/I subtype according to the degree of colocalization (highly/midly/lightly colocalized by the other subtype, corresponding to Division 1∼3). Each neuronal cell was assigned to one division based on its CCFv3 parcellations.^3^ The testing dataset was separated the same way. At the end, the divisions were combined into whole brain for cell neighborhood and class classification.

##### Data pre-processing

We observed that the image resolution of MERFISH imaging (167 nm^54^) is significantly higher than that of the rCCF image template. To avoid overlapping data, we adjusted the xy-plane image resolution of all images acquired in this study by upsampling each 2D plane by a factor of 10. We then removed the overlapping data (∼1.0% of the population) that retained the same xyz coordinates after image upsampling.

##### Data organization

Each training/testing dataset contained a table sized n × 21, where the n^th^ row corresponded to the n^th^ cell centroid; column #1∼14 corresponded to the values of fourteen spatially matched dMRI metrics; column #15∼21 corresponded to the z, x, y coordinates, cell class, cell neighborhood, neuron type, and brain division, respectively. In training by CCF structures, columns #1∼14 were substituted by the grayscale values of the AMBA’s STP image. Coordinates were included as predicting variables as they offered spatial information that is directly correlated to brain anatomy.^46^ Each predicting variable was rescaled to [0, 1] for data normalization. The cells with empty cell type labels, or outside of the brain, or mismatched during image registration, were deleted.

#### 5.5.3. Model architecture

We developed MFNet, a multi-diffusion-MRI fusion network for cell type prediction that integrates convolutional and transformer-based components (Figure 1D). The architecture consists of three modules—multi-scale convolutional feature extraction (Figure 1D, in green box), enhanced transformer encoding (Figure 1D, in orange box), and cross-modal feature fusion (Figure 1D, in red box)—followed by a classifier (Figure 1D, in blue box).

The convolutional module uses parallel 1D convolutions (kernels 3, 5, 7) to capture multi-scale features. Outputs are concatenated, refined with depthwise-separable and pointwise convolutions,^66^ and modulated by a channel-attention mechanism to enhance discriminative features.^67^ A residual connection stabilizes training.

The transformer module captures long-range dependencies^68^ by embedding inputs via convolutional projection with positional encodings,^69^ followed by a multi-layer transformer encoder. Global self-attention and a parallel local-attention branch extract complementary contextual features,^70^ which are combined via adaptive average pooling.

Cross-modal fusion aligns CNN and transformer^71^ features in a shared latent space using cross-attention, with a gating network adjusting their relative contributions.^72^ The fused representation is normalized and fed through fully connected layers with dropout to generate final cell type predictions.

#### 5.5.4. Model training and validation

The model was trained using a two-stage, three-level classification framework. In the first stage, for each predefined brain division under the excitatory/inhibitory (E/I) subtype scheme, the model classified whether a given cell belonged to the excitatory or inhibitory subtype. To address division-specific class imbalance, we defined loss-weight sets

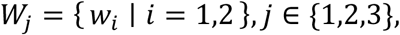

where 𝑤*_i_* represents the inverse prevalence of subtype 𝑖 within division 𝑗. Model optimization used a weighted cross-entropy loss,

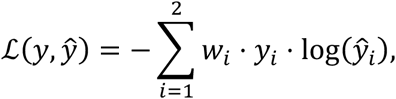

with 𝑦*_i_* ∈ {0,1} denoting the one-hot ground-truth label and *y_i_* the predicted probability for subtype 𝑖. This formulation increases the influence of underrepresented subtypes and compensates for imbalance across highly, moderately, and weakly co-localized divisions. Division-specific probability thresholds were estimated using held-out ground-truth data and subsequently applied during inference. To preserve anatomical consistency, predicted probabilities were further calibrated using a Z-slice spatial prior that adjusted scores according to each slice’s baseline E/I ratio prior to thresholding.

In the second stage, for all cells across the whole brain previously assigned an E/I subtype, the model simultaneously predicted each cell’s neighborhood and class.

The training process was conducted over 100 epochs for each division using an Adam optimizer with an initial learning rate of 1×10⁻⁴. To ensure reproducibility, a fixed random seed (42) was established across Python, NumPy, and PyTorch. All experiments were executed on an Intel I9-14900K CPU paired with an NVIDIA RTX 4090 GPU equipped with 24 GB of VRAM.

To validate the proposed model, we evaluated its performance by leave-p-groups-out validation using five independent slices from Section 5.5.2 as testing data. Classification performance was evaluated separately for the three classification levels. The classification metrics included weighted average accuracy and weighted average F1-score. Model selection relied solely on validation performance. We monitored training loss and validation accuracy throughout training. We used mild explicit regularization, including dropout with p = 0.3 in the classifier and L2 weight decay of 1 × 10^-5^, together with Batch Normalization.

#### 5.5.5. Ablation Study and Model Comparison Study

To measure the contribution of each module to the overall model, we ablated each component (Transformer, CNN, and fusion block, sequentially) on both neighborhood and class classification for excitatory and inhibitory neurons, and then the prediction accuracies after each ablation were recorded and compared. Meanwhile, we also compared the prediction accuracy on MFNet with MLP.

#### 5.5.6. UMAP projection

We performed a 2D UMAP projection to visualize the spatial clustering of neuron types, cell neighborhoods, and classes. To avoid memory issues, we down-sampled the complete dataset from Section 5.5.2 to a maximum of 1,000 cells per cell type class. We then used all predicting variables as input to create UMAP projections (n_neighbors=6, min_dist=0.5). The UMAPs projections were color-coded according to the corresponding cell types.

#### 5.5.7. Feature importance ranking

We ranked the predictive features based on their importance scores using the Analysis of Variance (ANOVA) filter. Features are ranked based on their F-statistics, enabling the selection of the most impactful ones. Feature ranking was done separately on neuron type, cell neighborhood, and cell class classifications.

#### 5.5.8. Statistical analysis

All statistical analyses were performed using GraphPad Prism version 10.6.1 (GraphPad Software, Inc.). Paired comparisons between conditions were evaluated using the nonparametric Wilcoxon matched-pairs signed-rank test. Multiple-comparison adjustment by Benjamini-Hochberg for false-discovery-rate (FDR) correction was applied, and adjusted p < 0.05 was considered statistically significant.

### 5.6. MRI correlation tests with spatial transcriptomics and spatial autocorrelation correction

To examine the correlation between MRI and spatial transcriptomics, we performed a linear correlation test on the fourteen CCF-registered dMRI metrics and the imputed MERFISH spatial transcriptomic data of NT type marker genes (or MERFISH spatial transcriptomic data of 500 genes). The imputed MERFISH spatial transcriptomics were obtained from Yao et al.^3^ Regions of interest (ROIs, *n* = 686) were selected as all voxels within each of the CCFv3 parcellation, and the mean value for each ROI was used as one data point in the correlation test (Figure 1E). We recorded all the pairwise Pearson’s correlation coefficients (PCC) as observed PCCs.

Spatial autocorrelation (SA) in dMRI maps was corrected using BrainSMASH^73^ to generate spatially autocorrelated surrogate maps (n = 1000 per dMRI map) whose spatial dependence matched that of the dMRI data (Figure S9). For each gene, we calculated the PCCs between its expression and these surrogate maps as the surrogate PCCs, forming a null distribution. For each gene-dMRI pair, the percentage of surrogate PCCs exceeding the observed PCC was calculated as the SA-corrected p-value. SA-corrected p-value < 0.05 was considered statistically significant.

### 5.7. Complete whole-brain cell type atlas at CCFv3 space generation, validation, and evaluation

Using the MFNet, we generated a complete whole-brain cell type atlas (MFNet Atlas, Figure 1F) in CCFv3 space. First, we applied the pre-processed and CCFv3-registered neuron cell segmentation result images from 3D NeuN images to spatially matched dMRI maps to filter the neuronal cell voxels. Next, we predicted firstly the E/I neuron type and then the cell neighborhoods and classes on these voxels. The MFNet-generated 3D whole-brain neuronal cell atlas, including neuron type, cell neighborhood, and class annotations, was saved as NIFTI files and made available through our data portal (https://github.com/selinahxy/mricelltype). The same color scheme as the ABC Atlas was used. Lastly, to evaluate the MFNet Atlas, we compared the similarity between the MFNet Atlas and the ABC Atlas by using histogram correlation metrics between pairwise spatially matched coronal slices.

To examine the advantages of the MFNet Atlas over the ABC Atlas, we first interpolated the interval z-slices in between the available ABC Atlas z-slices by using k-nearest-neighbor interpolation across z. For each target slice, we locate the k closest ABC slices in z, weight them by inverse distance, and propagate x–y coordinates and labels via distance-weighted majority vote; ties follow the empirical class prior. At the boundaries, we clamp to the nearest observed slice. This yields a dense 1320-slice atlas without altering in-plane geometry or applying smoothing. Then we compared the inhibitory neuron type maps of MFNet Alas with the interpolated ABC Atlas, with Gad1 and Gad2 ISH images as the ground truth.

## Acknowledgments

This work was funded by the National Institutes of Health grants R01 NS125020 (N.W.), P41EB015897 (G.A.J), S10OD010683 (G.A.J). The funders had no role in study design, data collection and analysis, decision to publish, or preparation of the manuscript.

## Ethical Approval

All animal studies have been approved by the appropriate ethics committee: Duke University Institutional Animal Care and Use Committee. Approval code A226-17-09.

## Declaration of Interest

The authors declare no competing interests.

## Data Availability Statement

The data and codes that support the findings of this study are available from the data sharing repository: https://utsw.box.com/s/mhjwdck148p3y3vofd22j7pd0h5jdhby and the GitHub repository: https://github.com/selinahxy/mricelltype. Availabilities about publicly available datasets can be found in Table S1.

## Declaration of generative AI and AI-assisted technologies in the manuscript preparation process

During the preparation of this work the author(s) used ChatGPT (GPT-5.1) in order to improve charity and grammar. After using this tool/service, the authors reviewed and edited the content as needed and take full responsibility for the content of the published article.

## Notes

https://utsw.box.com/s/mhjwdck148p3y3vofd22j7pd0h5jdhby

https://github.com/selinahxy/mricelltype

### Competing Interest Statement

The authors have declared no competing interest.

