## Supplemental Materials for "High-resolution MRI Guided Whole Mouse Brain Cell Type Atlas using Deep Learning"

### Supplementary Materials

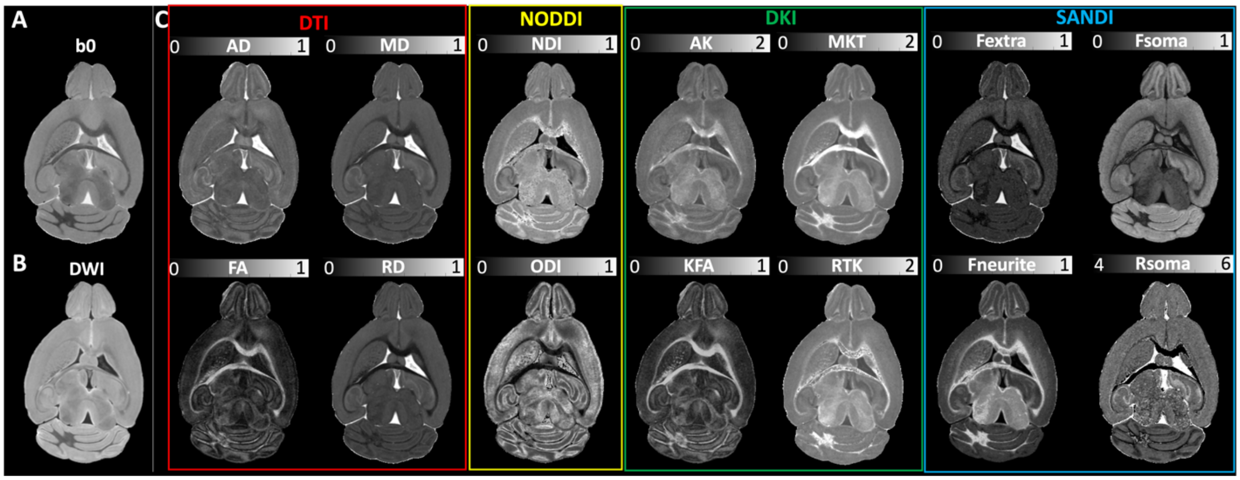


*Figure S1. MRI images at native space. (A) b0 image. (B) Diffusion-weighted image. (C) dMRI maps. dMRI maps include parametric maps from DTI (in red box), NODDI (in yellow box), DKI (in green box), and SANDI (in blue box) models.*

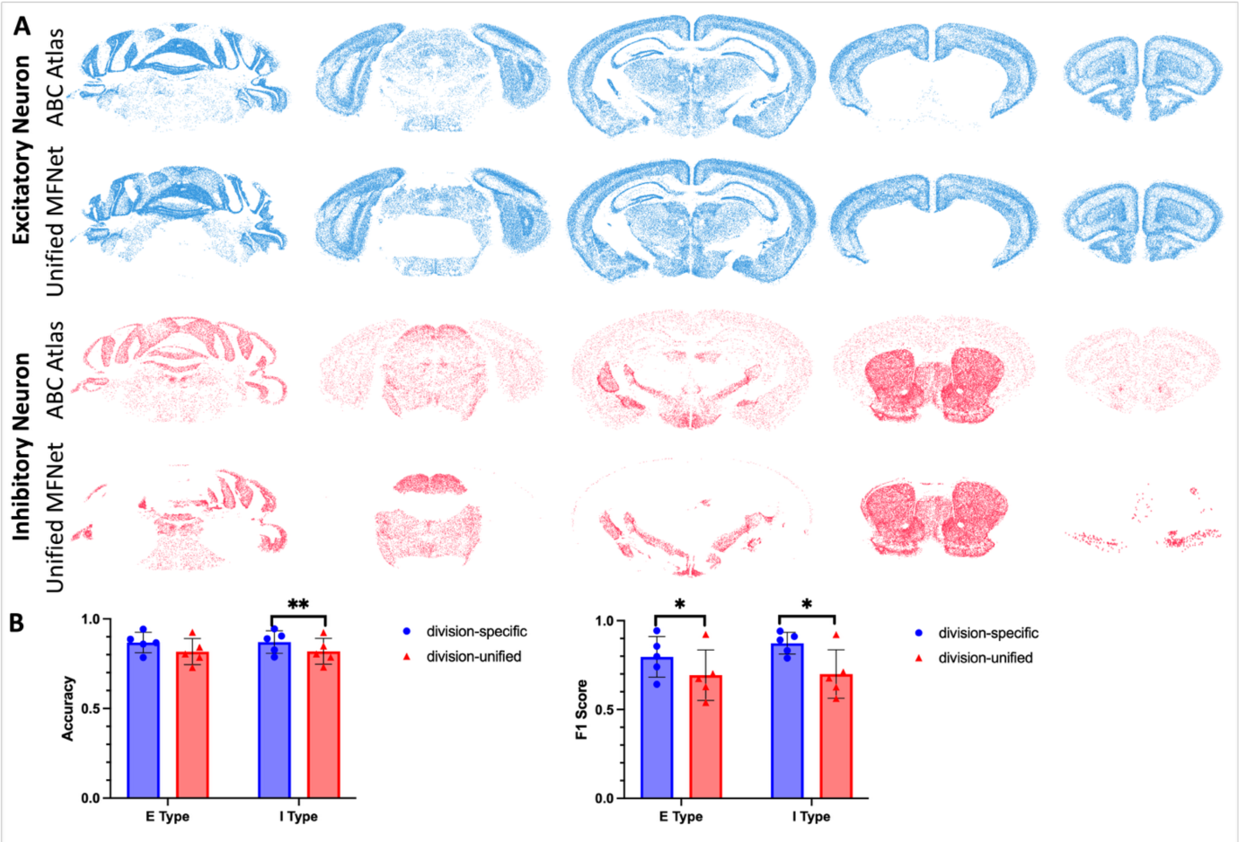


*Figure S2. Neuronal cell type prediction using division-unified MFNet. (A) Prediction results on neuron types in the mouse whole brain from each one of the testing slices. From top to bottom: ABC Atlas and division-unified MFNet*

*predicted. (B) Bar graph comparing the weighted average prediction accuracy (left) and F1 score (right) in by division-specific MFNet (in blue bars) and division-unified MFNet (in red bars). Error bars represent the standard deviation among the testing slices. n = 5. * FDR-corrected p-values < 0.05, ** FDR-corrected p-values < 0.01.*

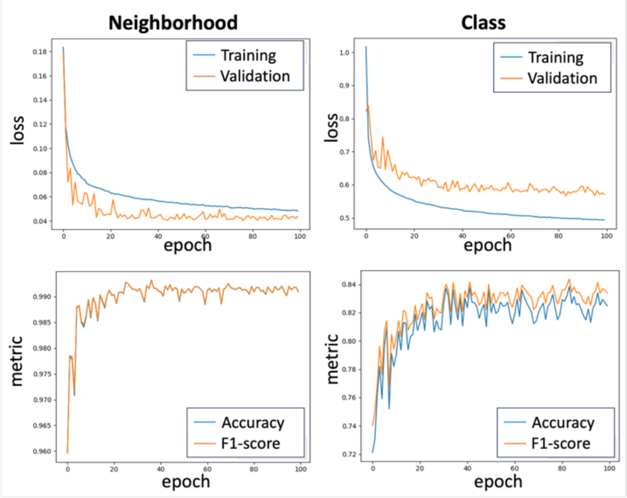


*Figure S3. Loss curves (top) during training and validation, and metric curves (bottom) during validation.*

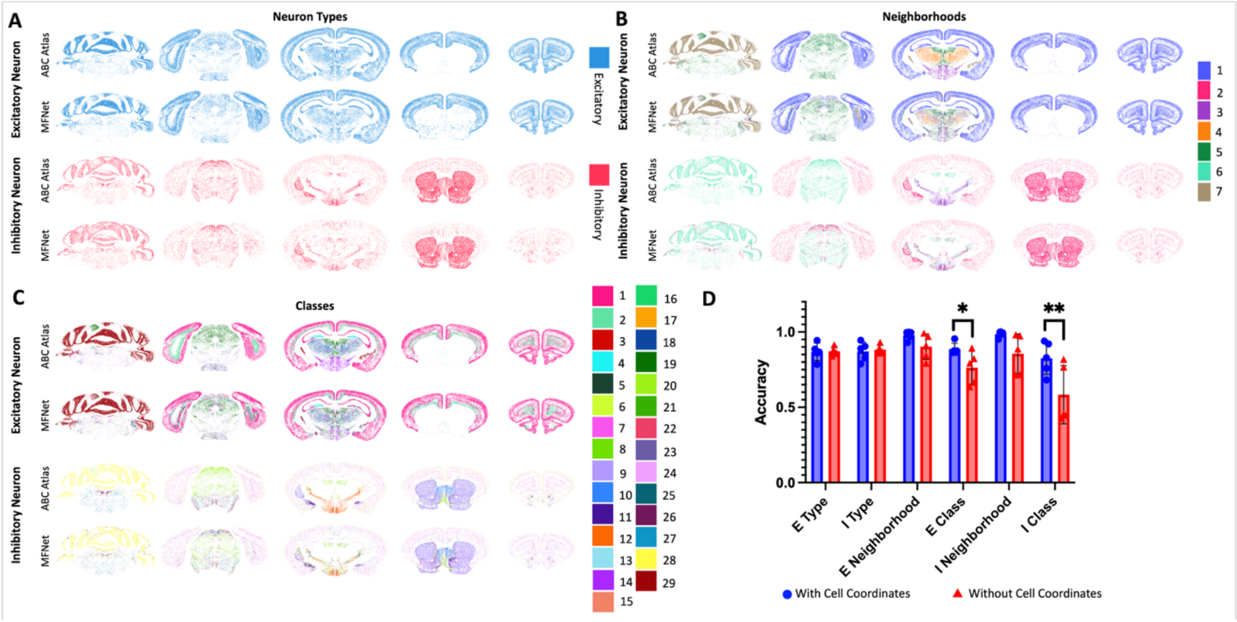


*Figure S4. Comparison on neuronal cell type prediction with or without cell coordinates. (A~C) Prediction results on neuron types (A), neuronal cell neighborhoods (B) and classes (C) in the mouse whole brain from each one of the testing slices. From top to bottom: excitatory neurons from ABC Atlas and MFNet predicted, inhibitory neurons from ABC Atlas and MFNet predicted. (D) Bar graph comparing the weighted average prediction accuracy with (in blue bars) and without cell coordinates (in red bars). Error bars represent the standard deviation among the testing slices. n = 5. * FDR-corrected p-values < 0.05. ** FDR-corrected p-values < 0.01.*

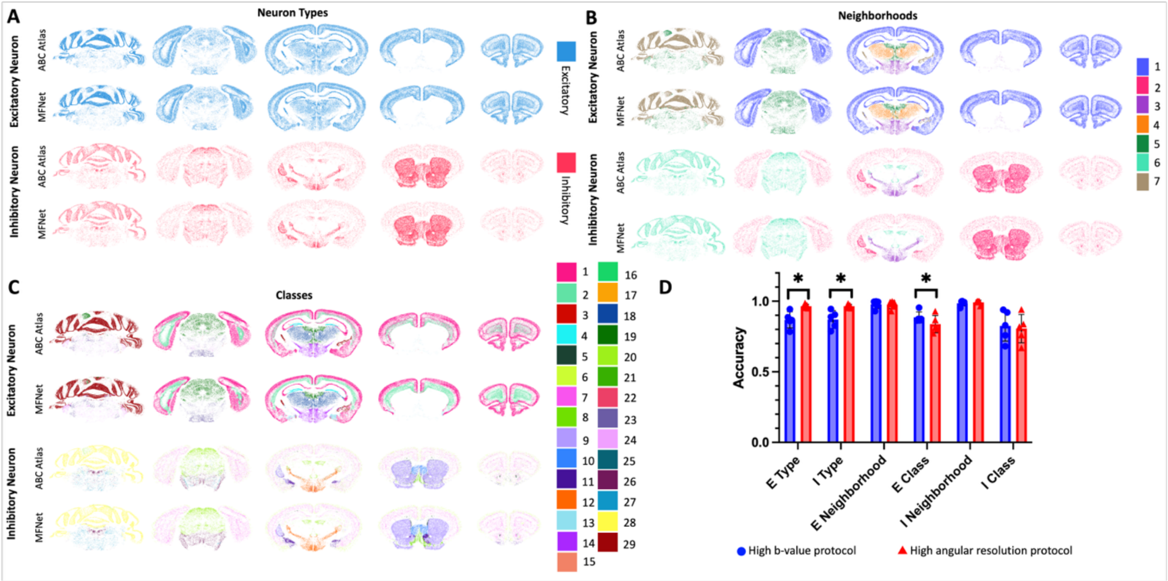


*Figure S5. Neuronal cell type prediction using MRI data from high angular resolution protocol. (A~C) Prediction results on neuron types (A), neuronal cell neighborhoods (B) and classes (C) in the mouse whole brain from each one of the testing slices. From top to bottom: excitatory neurons from ABC Atlas and MFNet predicted, inhibitory neurons from ABC Atlas and MFNet predicted. (D) Bar graph comparing the weighted average prediction accuracy by high b-value protocol (in blue bars) and high angular resolution protocol (in red bars). Error bars represent the standard deviation among the testing slices. n = 5. * FDR-corrected p-values < 0.05.*

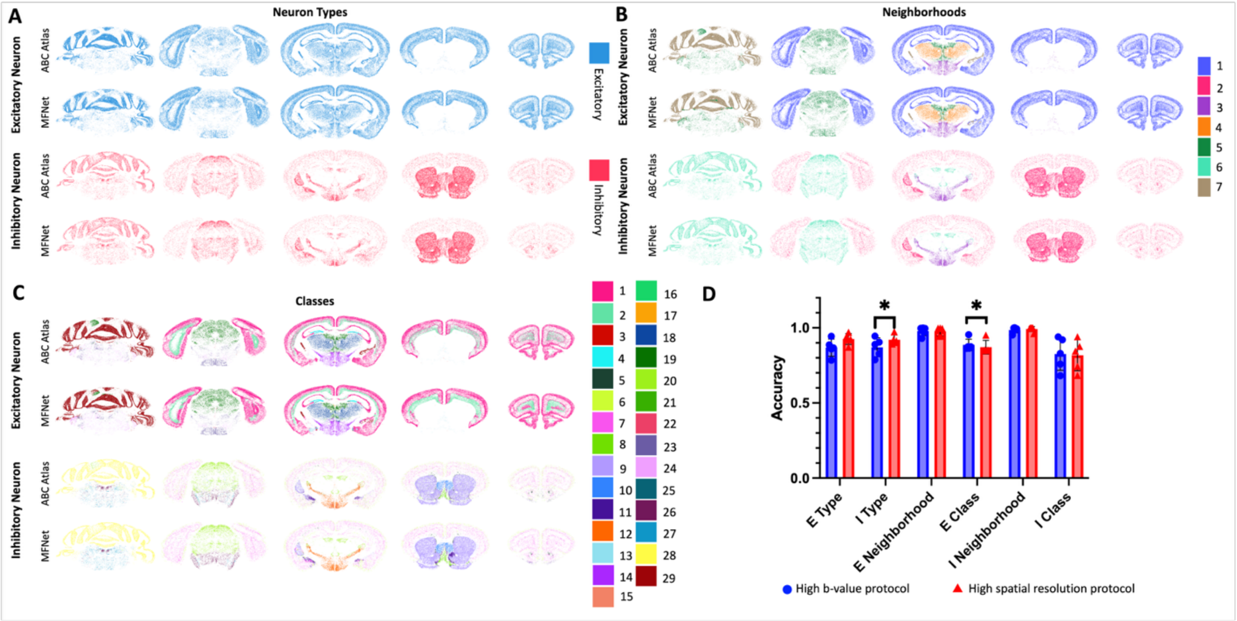


*Figure S6. Comparison on neuronal cell type prediction with MRI data from high spatial resolution protocol. (A~C) Prediction results on neuron types (A), neuronal cell neighborhoods (B) and classes (C) in the mouse whole brain from each one of the testing slices. From top to bottom: excitatory neurons from ABC Atlas and MFNet predicted, inhibitory neurons from ABC Atlas and MFNet predicted. (D) Bar graph comparing the weighted average prediction*

*accuracy by high b-value protocol (in blue bars) and high spatial resolution protocol (in red bars). Error bars represent the standard deviation among the testing slices. n = 5. * FDR-corrected p-values < 0.05.*

.

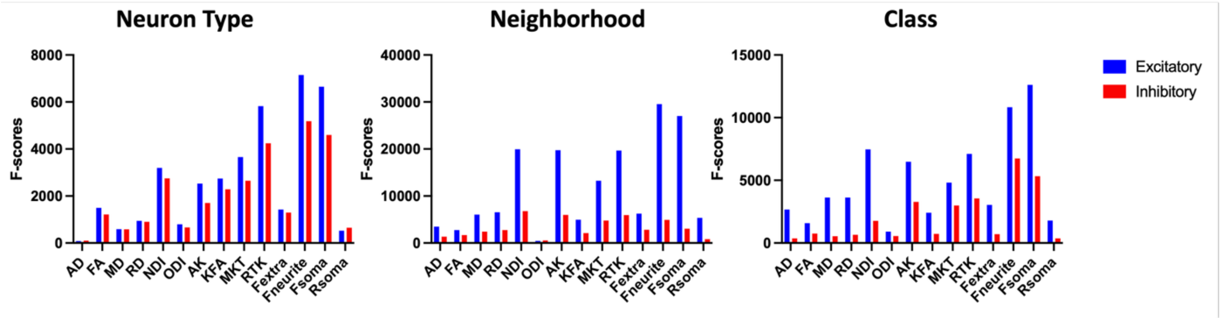


*Figure S7. Predictive feature importance on predicting neuron type (left), neighborhood (middle), and class (right). F-scores are shown separately for excitatory (in blue bars) and inhibitory (in red bars) neuron predictions.*

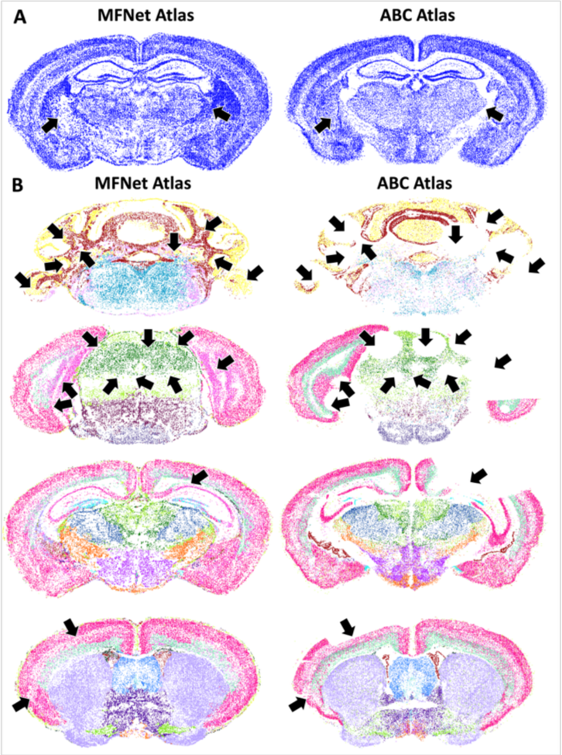


*Figure S8. (A) Comparisons between one representative slice from MFNet Atlas (left) and corresponding ABC Atlas (right). The two atlases are color-coded by neuron cell segmentation. Arrows indicate the major miss-matched areas (B) Comparisons between two representative slices from MFNet Atlas (left) and corresponding ABC Atlas (right) that were broken. The two atlases are color-coded by cell class of all neurons. Arrows indicate the broken areas.*

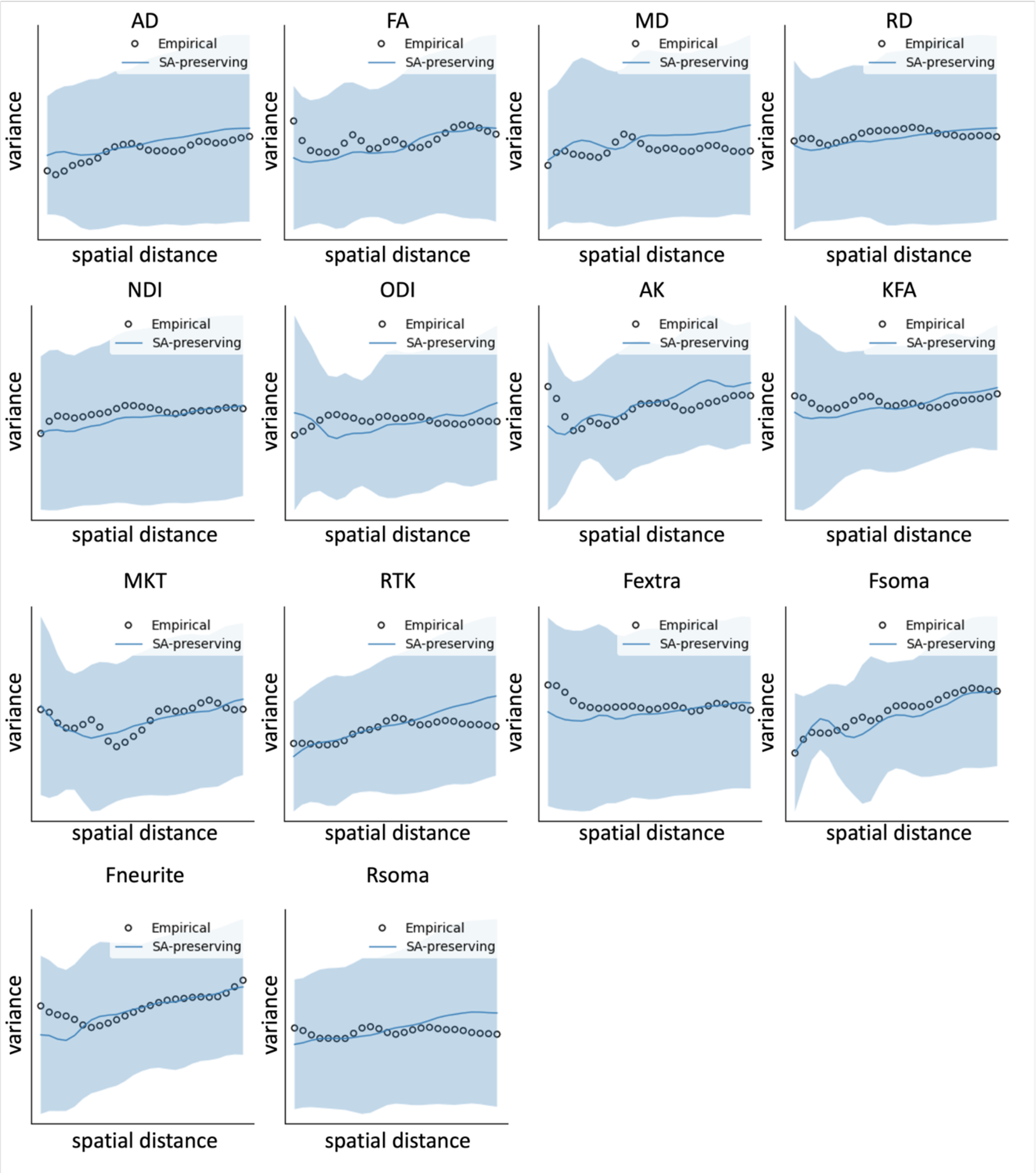


*Figure S9. Variograms of empirical dMRI maps and spatial autocorrelation (SA)-preserving surrogate maps. Black circle dots = empirical dMRI map, blue line = SA-preserving surrogate maps.*

Table S1. Multi-modality atlas data description

| Data Name | Data source | References | Description |
| --- | --- | --- | --- |
| dMRI Images | https://utsw.box.com/s/mhjwdck148p3y3vofd22j7pd0h5jdhby | In-house imaged | dMRI images including metrics from DTI, NODDI, DKI, and SANDI models covering the whole mouse brain of B6 and 5xFAD mice |
| ABA Template | https://portal.brain-map.org/atlases-and-data/bkp/abc-atlas | [1] | The Allen Mouse Brain Atlas (AMBA)'s two-photon tomography image templates in Allen Mouse Brain Common Coordinate Framework 10 um (CCFv3) and MERFISH CCF mapped coordinates (resampled CCF) |
| ABA Annotation | https://portal.brain-map.org/atlases-and-data/bkp/abc-atlas | [1] | The AMBA's 3D annotation labels in CCFv3 and resampled CCF |
| 3D NeuN Image | https://civmimagespace.civm.duhs.duke.edu/ | [2] | Stains neuronal nuclei in the central nervous systems in whole mouse brain |
| 2D GAD1 ISH Image | https://mouse.brain-map.org/ | [1] | In situ hybridization (ISH) image of Gad1 expression through the |

| | | | |
|---|---|---|---|
| | | | adult mouse brain |
| 2D GAD2 ISH Image | https://mouse.brain-map.org/ | [1] | ISH image of Gad2 expression through the adult mouse brain |

Table S2. Ablation study and model comparison results

| | | MFNet | W/O Transformer | W/O CNN | W/O Fusion Block | MLP |
|---|---|---|---|---|---|---|
| Inhibitory Neuron | Class | 99.32 | 98.71 | 96.84 | 98.95 | 97.92 |
| | Neighborhood | 83.84 | 82.69 | 81.21 | 83.26 | 82.34 |
| Excitatory Neuron | Class | 97.13 | 96.28 | 94.51 | 96.61 | 95.57 |
| | Neighborhood | 85.23 | 84.37 | 82.46 | 84.71 | 83.62 |
